# SINHCAF/FAM60A links SIN3A function to the hypoxia response and its levels are predictive of cancer patient survival

**DOI:** 10.1101/176032

**Authors:** John Biddlestone, Michael Batie, Alena Shmakova, Daniel Bandarra, Elena V. Knatko, Albena T. Dinkova-Kostova, Ivan Munoz, Ramasubramanian Sundaramoorthy, Tom Owen-Hughes, Sonia Rocha

**Author notes:** Telephone for corresponding author (*): Sonia Rocha. e-mail addresses of all authors: John Biddlestone Michael Batie Alena Shmakova Daniel Bandarra Elena Knatko Albena Dinkova-Kostova Ivan Munoz Ramasubramanian Sundaramoorthy Tom Owen-Hughes Sonia Rocha.

## Abstract

The SIN3A-HDAC complex is a master transcriptional repressor, required for development but often deregulated in disease. Here, we report that the recently identified new component of this complex, SINHCAF/FAM60A, links the SIN3A-HDAC co-repressor complex function to the hypoxia response. SINHCAF Chromatin Immunoprecipitation-sequencing and gene expression analysis reveal a signature associated with the activation of the hypoxia response. We show that SINHCAF specifically repress HIF 2α mRNA and protein expression resulting in functional cellular changes in *in-vitro* angiogenesis, and proliferation. Analysis of patient datasets demonstrates that SINHCAF and HIF 2α mRNA levels are inversely correlated and predict contrasting outcomes for patient survival in both colon and lung cancer. This relationship is also observed in a mouse model of colon cancer, indicating an evolutionary conserved mechanism. Our analysis reveals an unexpected link between SINHCAF and cancer cell signalling via regulation of the hypoxia response that is predictive of poor patient outcome.

## Introduction

SINHCAF/FAM60A is a poorly characterised protein that has been shown to be part of the SIN3A repressor complex by two independent studies (Munoz et al., 2012; Smith et al., 2012). SIN3A is a multi-protein complex with function in developmental processes such as stem cell function (Saunders et al., 2017) but also in pathological processes such as cancer (Kadamb et al., 2013). It controls cellular metabolism, cell cycle and cell survival (Kadamb et al., 2013). More recently, it a single nucleotide polymorphism on SINHCAF has been associated with Type II diabetes in a cohort of Japanese patients (Imamura et al., 2016). Finally, it has been associated with esophageal cancer, by an integrative analysis of copy number (Dong et al., 2017).

Solid tumours are often characterised by the presence of low oxygen or hypoxia (Moniz et al., 2014). Adaptation and survival to such environment is mediated by a family of transcription factors called Hypoxia Inducible Factors (HIFs). Transcriptional target genes that are controlled by the HIFs, code for proteins that are involved in important cellular processes including energy homeostasis, migration, and differentiation (Keith and Simon, 2007; Rocha, 2007; Schofield and Ratcliffe, 2004). Deregulation of the HIF system has been shown to be important in the development of multiple disease processes including cancer progression and stem cell differentiation (Semenza, 2008, 2012).

HIF is a heterodimeric transcription factor that consists of a constitutively expressed HIF 1β subunit and an O_2_-regulated HIF-α subunit (Wang et al., 1995). Three isoforms of HIF α have been identified (HIF 1α, 2α, and 3α). The HIF-α isoforms are all characterized by the presence of bHLH (basic helix–loop–helix)–PAS (Per/ARNT/Sim), and ODD (oxygen-dependent degradation) domains. Both HIF-1α and HIF 2α have important cellular functions as transcription factors with some redundancy in their targets (Carroll and Ashcroft, 2006; Hu et al., 2006). HIF-2α protein shares sequence similarity and functional overlap with HIF-1α, but its distribution is restricted to certain cell types, and in some cases, it mediates distinct biological functions (Patel and Simon, 2008).

The regulation of the HIF α subunits is best understood at the post-transcriptional level, and is mediated by hydroxylation-dependent proteasomal degradation. In well-oxygenated cells, HIF-α is hydroxylated in its ODD. This proline hydroxylation is catalysed by a class of dioxygenase enzymes called prolyl hydroxylases (PHDs). PHDs require Fe^2+^ and α-ketoglutarate (α-KG) as co-factors for their catalytic activity and have an absolute requirement for molecular oxygen as a co-substrate, making their activity reduced in hypoxia (Bruegge et al., 2007; Epstein et al., 2001; Fandrey et al., 2006; Frede et al., 2006). Prolyl-hydroxylation of HIF α attracts the von Hippel-Lindau (vHL) tumor suppressor protein, which recruits the Elongin C-Elongin B-Cullin 2-E3-ubiquitin-ligase complex, leading to the Lys48-linked poly-ubiquitination and proteasomal degradation of HIF α (Ivan et al., 2001; Jaakkola et al., 2001; Yu et al., 2001). In hypoxia, HIF α is stabilized, can form a heterodimer with HIF 1β in the nucleus and bind to the consensus cis-acting hypoxia response element (HRE) nucleotide sequence 5’-RCGTG-3’, which is present within the enhancers and/or promoters of HIF target genes (Pugh et al., 1991; Schodel et al., 2011; Semenza et al., 1996). HIF α stabilisation therefore allows the cell to enact a transcriptional programme that is appropriate to the hypoxic environment (Kaelin and Ratcliffe, 2008).

In contrast to the post-translation regulation of HIF, the factors that regulate the basal expression of the HIF α isoforms are only now being investigated. Deregulation of HIF basal expression has been linked to the development of solid tumours (Kazi et al., 2014). The transcriptional regulator NF-κB is a direct modulator of HIF-1α expression, in basal and hypoxic conditions, as well as in response to inflammatory stimulus (Belaiba et al., 2007; Bonello et al., 2007; Frede et al., 2006; Rius et al., 2008; van Uden et al., 2008). NF-κB also directly regulates HIF-1β mRNA and protein levels, resulting in modulation of HIF 2α protein levels by preventing protein degradation(Bandarra et al., 2014; van Uden et al., 2011). Additional studies have also shown that SP1/3 and Egr-1 transcription factors and the STAT3 transcriptional activator can all regulate expression of HIF 1α RNA (Biswas et al., 2013; Koshikawa et al., 2009; Lafleur et al., 2014; Niu et al., 2008; Oh et al., 2008; Patel and Kalra, 2010). HIF 2α is regulated by SP1/3 (Wada et al., 2006) and during cell progression by E2F1 (Moniz et al., 2015). Interestingly, HIF 2α has recently been shown to be sensitive to Histone Deacetylase (HDAC) inhibition in soft tissue sarcomas (Nakazawa et al., 2016). HDACs are components of several transcriptional repressor complexes, including the SIN3A complex (Kadamb et al., 2013). SINHCAF (SIN3A-HDAC associated factor/FAM60A) is a relatively new component of the SIN3A-HDAC complex (Munoz et al., 2012; Smith et al., 2012; Streubel et al., 2017). It has been associated with regulation if Cyclin D1 (Munoz et al., 2012), TGF-β signalling (Smith et al., 2012) and more recently stem cell maintenance (Streubel et al., 2017).

Here, we show that SINHCAF links SIN3A function to hypoxia signalling. ChIP-sequencing analysis for SINHCAF and integrative analysis for SIN3A, identify a hypoxia specific gene signature. SINHCAF targets the HIF 2α promoter, resulting in histone deacetylation and gene silencing. Analysis of cancer transcript datasets reveals that SINHCAF and HIF 2α expression is inversely correlated and predicts contrasting patient survival outcomes in colon and lung cancers, an effect also observed in a mouse model of colon cancer. Thus, SINHCAF links SIN3A and hypoxia signalling and is associated with poor patient survival outcome.

## Results

### ChIP-sequencing analysis of SINHCAF reveals its function as a transcriptional regulator

SINHCAF has been identified as a component of the SIN3A/HDAC complex (Munoz et al., 2012; Smith et al., 2012), and to contribute to transcriptional control of Cyclin D1 (Munoz et al., 2012). Microarray analysis of SINHCAF and SIN3A regulated genes in A549 cells has identified co-regulated genes within the TGF-β signalling pathway (Smith et al., 2012). However, it is currently unknown if and how SINHCAF correlates with SIN3A functionally. To identify direct targets of SINHCAF, we performed ChIP-sequencing analysis for endogenous SINHCAF, using two different cellular backgrounds, HeLa a human cervical carcinoma cell line and DLD-1 a human colorectal carcinoma cell line. Analysis of the sequencing data revealed a significant overlap between the peaks observed in HeLa and DLD-1 cells, with around 60% of the peaks being identified in both cellular backgrounds (Figure 1A). Mapping of the sequencing reads to the genome architecture revealed that 93% of the identified peaks in both HeLa and DLD-1 cells were present within 3000 bp of transcriptional start sites (Figure 1B, Sup. Figure S1A), with the remaining being located in the gene body (3%) and in intergenic regions (4%). This suggests that SINHCAF is a transcriptional regulator. To further characterise the location of SINHCAF across the genome, we plotted all the shared peaks with respect to the transcriptional start site (TSS). This analysis demonstrated that SINHCAF is mainly located at the TSS (Figure 1C; Sup. Figure S1B), further suggesting its role as a transcriptional regulator.

**Figure 1:**
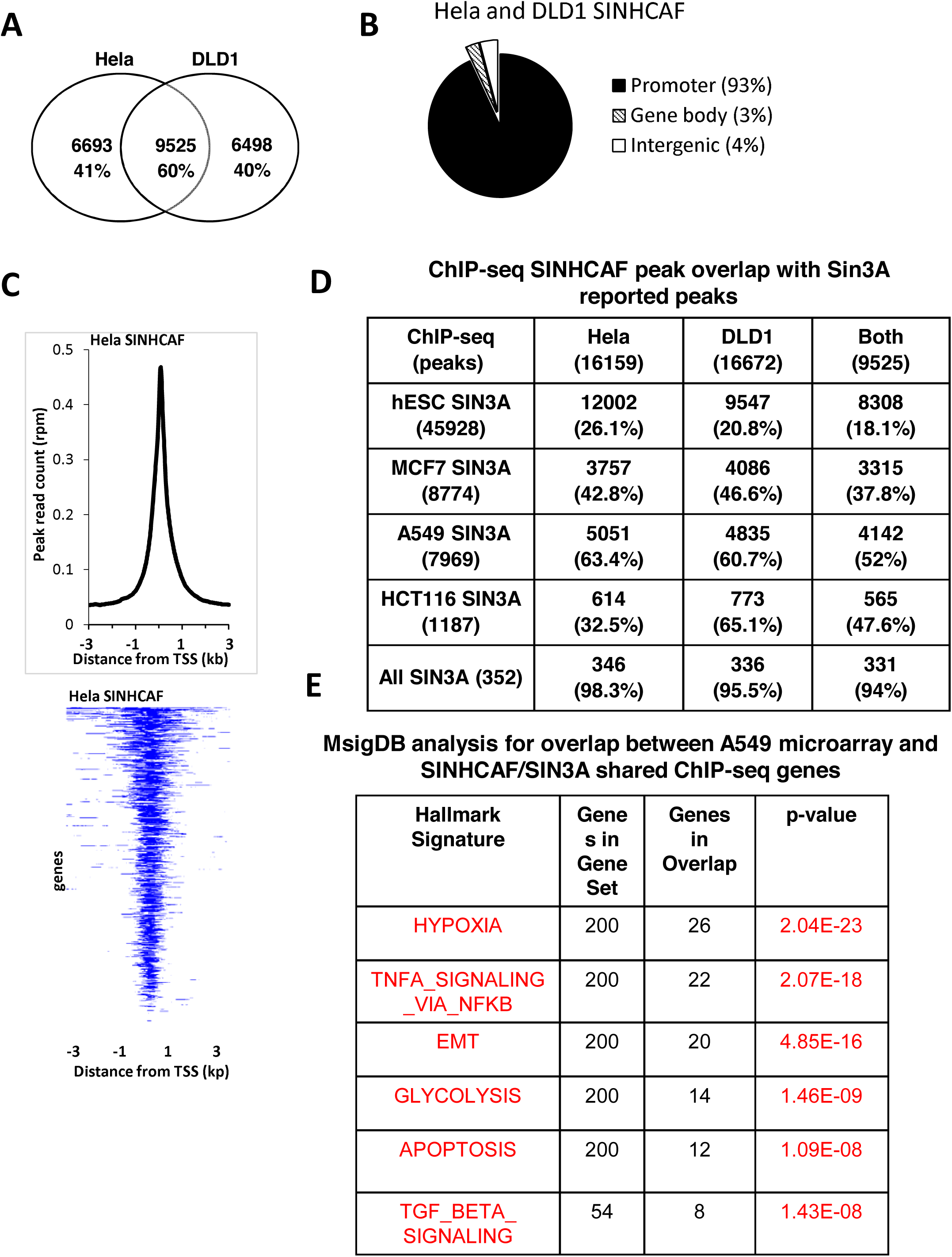
SINHCAF is associated with gene promoters and links SIN3A function to Hypoxia and Inflammation. (A) Venn diagram depicting number of peak overlap between SINHCAF ChIP-sequencing analysis in two distinct cellular backgrounds, HeLa and DLD-1. (B) Genomic distribution for SINHCAF shared peaks between HeLa and DLD-1. (C) Annotation of HeLa SINHCAF ChIP-sequencing peaks in reference to the TSS. (D) Analysis of SINHCAF ChIP-sequening peak overlap with published datasets for SIN3A ChIP-sequencing across different cellular backgrounds. (E) Molecular signature analysis using GSEA for the overlap between shared SINHCAF/SIN3A genes and A540 microarray following SINHCAF depletion. See also Figure S1.

Given SINHCAF’s known association with SIN3A, we investigated how SINHCAF location overlapped with published ChIP-sequencing datasets for SIN3A (Figure 1D). We analysed 4 distinct SIN3A ChIP-sequencing datasets across 4 different cellular backgrounds. Percentage of overlap varied across cell lines as expected, from the lowest 20.8% to the highest of 65.1% (Figure 1D). This variation was due to the intrinsic nature of the number of peaks called in each of the cell types analysed. Interestingly, when we created a list of SIN3A peaks observed across all cellular backgrounds and datasets, the overlap with SINHCAF peaks rose to a minimal of 94%, demonstrating that indeed, SINHCAF is part of the core and conserved function of SIN3A complex across cell types (Figure 1D). Furthermore, this data demonstrated that SIN3A function is cell type specific, since only 0.8 to maximum 29.7% of all identified SIN3A occupied genes are conserved across different cellular backgrounds (Figure 1D).

To investigate the function of SINHCAF in transcriptional regulation at these bound sites we made use of published data generated in the lung adenocarcinoma cell line A549 (Smith et al., 2012). We performed gene set enrichment analysis (GSEA) (Subramanian et al., 2005) on the overlap on the genes that are both regulated at the transcript level by loss of SINHCAF and bound by SINHCAF and SIN3A in ChIP-sequencing datasets (Figure 1E, Sup. Table 1). While we did find an enrichment for TGF-β signalling genes in this analysis, the top hallmark signalling pathways were “Hypoxia” and “TNF-α signalling via NF-κB” (Figure 1E). While HDAC function has been associated with both hypoxia and inflammation, no association of SIN3A function has been reported with both Hypoxia or NF-κB mediated signalling pathways. However, we had previously demonstrated that hypoxia and NF-κB are part of an intricate crosstalk (Biddlestone et al., 2015; D’Ignazio et al., 2016; D’Ignazio et al., 2017).

**Table 1:**
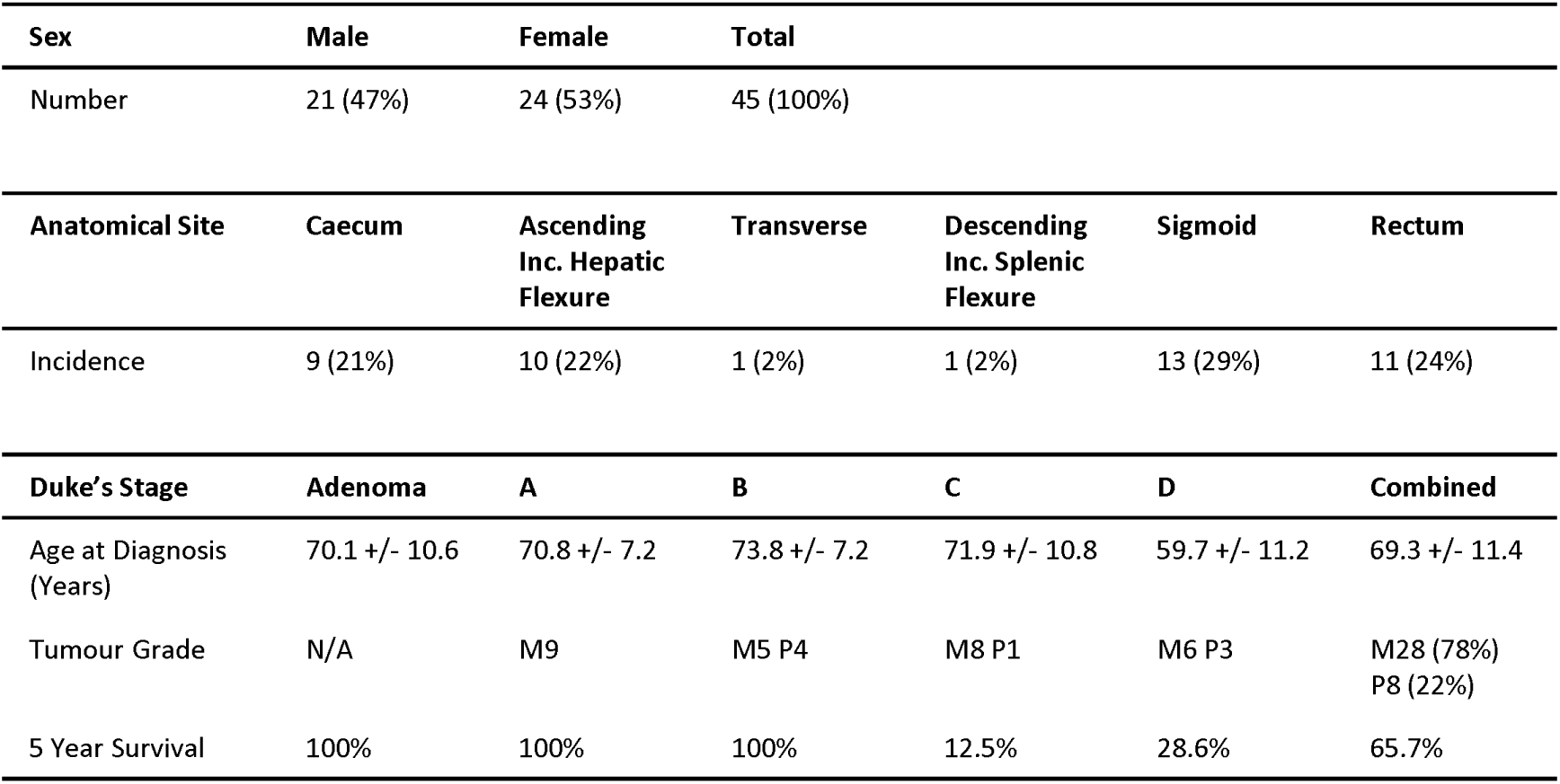
Colorectal Cancer Patient Demographics. Distribution of several demographics for patients whose fresh tissue was used to extract RNA for analysis of the contribution of SINHCAF/HIF-2α to colorectal cancer. Controls were paired samples of healthy tissue from same patients. Data shown are in format mean +/ S.D.

### SINHCAF is a repressor of HIF 2α

Analysis of the hypoxia related genes identified by GSEA in Figure 1E, revealed that HIF 2α (gene name *EPAS1*) is a gene repressed by both SINHCAF and SIN3A (Figure 2A-B). Very little is known about the regulation of the HIF 2α gene, apart that it can be regulated by E2F1 (Moniz et al., 2015), its promoter can be methylated (Cruzeiro et al., 2017; Cui et al., 2016), and that HDACs can repress both HIF 2α and HIF 1α genes (Nakazawa et al., 2016).

**Figure 2:**
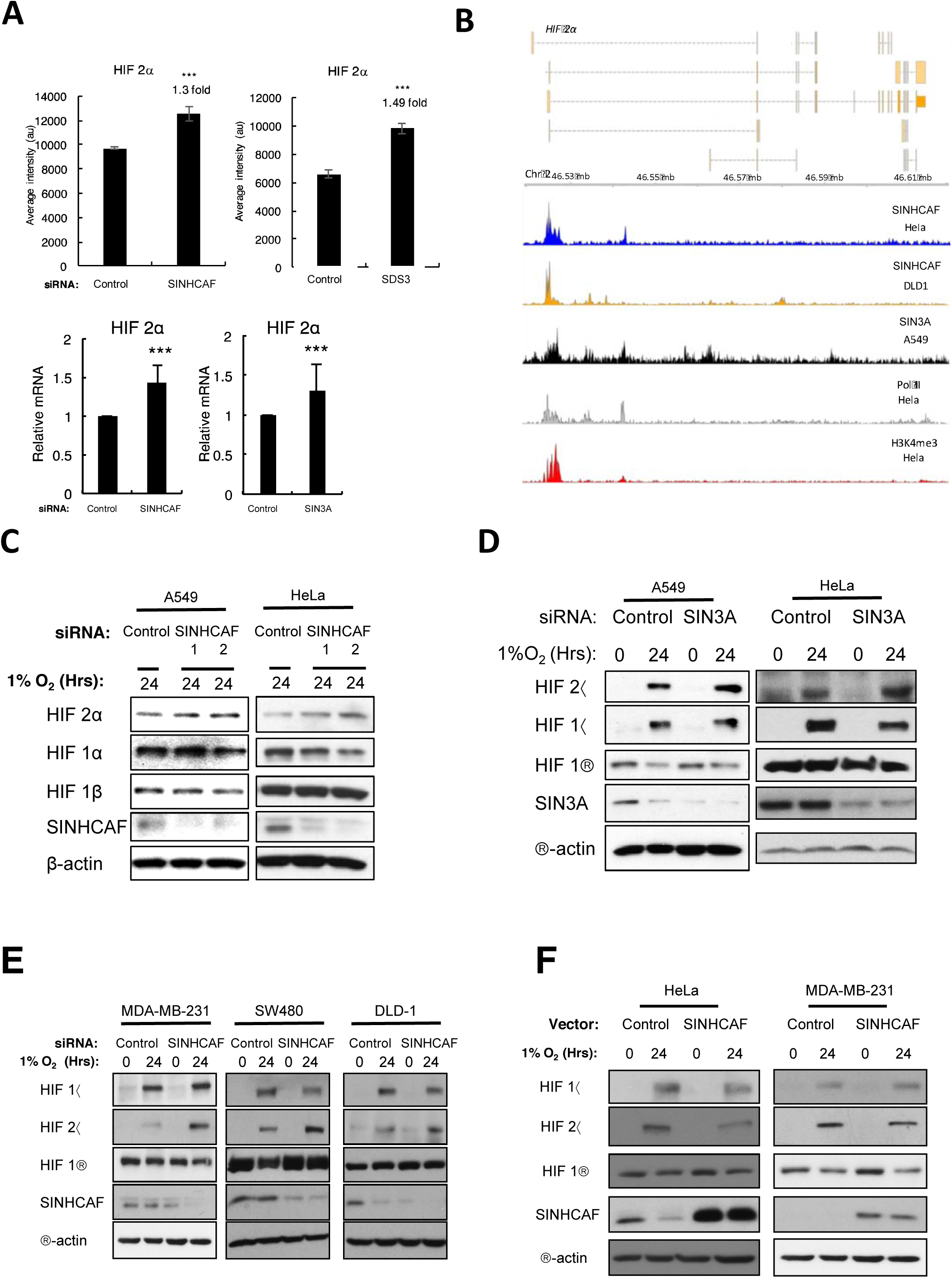
SINHCAF is a Repressor of HIF 2α in Multiple Cell Lines. (A) Microarray analysis levels for HIF 2α following SINHCAF and SDS3 in A549 cells (top panel). qPCR validation using A549 cells, and independent sets of siRNA oligonucleotides for SINHCAF and SIN3A (lower panel). Graphs depict mean + S.E.M. * P ≤ 0.05, ** P ≤ 0.01, *** P ≤ 0.001. (B) Gene track analysis of SINHCAF ChIP-sequencing in HeLa and DLD-1 cells, with overlap with SIN3A ChIP-sequencing in A549 cells. Pol II and H3K4me3 peaks for HeLa are also shown. (C) Control or one of two SINHCAF [1/2] siRNA oligonucleotides were transfected to A549 and HeLa cells cultured in the presence of hypoxia for 24 hours. Lysed samples were analysed by immunoblot for expression of HIF system isoforms and SINHCAF. (D) Control or SIN3A siRNA oligonucleotides were transfected to A549 and HeLa cells cultured in normoxia or hypoxia for 24 hours. Lysed samples were analysed by immunoblot for expression of HIF system isoforms and SIN3A. (E) Expression of HIF 2α following knockdown of SINHCAF and exposure to hypoxia for 24 hours was determined in breast MDA-MB-231 and two colorectal (SW480, DLD-1) cancer cell lines. (D) SINHCAF was overexpressed in HeLa and MDA-MB-231 cells with or without exposure to hypoxia for 24 hours. Lysed samples were analysed by immunoblot for expression of HIF system isoforms and SINHCAF. Representative images from at least three experiments are shown.

To determine if SINHCAF is a novel regulator of HIF 2α, A549 and HeLa cells were transfected with control or two SINHCAF siRNA oligonucleotides in order to examine the effect of SINHCAF knockdown on the expression of the HIF isoforms (Figure 2C). Knockdown of SINHCAF resulted in an increase in HIF 2α, but not HIF 1α or HIF 1β protein expression following exposure to hypoxia for 24 hours. Similar results were also observed when SIN3A was depleted by siRNA, with elevated levels of HIF 2α occurring in both HeLa and A549 cells (Figure 2D). To determine the penetrance of this effect similar experiments were performed in multiple cell lines. Loss of SINHCAF resulted in significant increases in HIF 2α with little or no change to HIF-1α protein following exposure to hypoxia in breast cancer cells (MDA-MB-231), and two colorectal (SW480, DLD-1) cell lines (Figure 2E). In addition, overexpression of control or SINHCAF DNA plasmids in cells was performed to determine if gain of function experiments would lead to the opposite effect on HIF 2α levels. Overexpression of SINHCAF resulted in a significant decrease in HIF 2α protein following exposure to hypoxia for 24 hours in both HeLa and MDA-MB-231 cells confirming that the siRNA results are not a technical artefact but also the responsiveness of the system (Figure 2F).

### SINHCAF regulates the HIF 2α promoter directly and is important for HDAC1 recruitment

The study of the regulation HIF α proteins has been mainly characterised by a control at the post-transcriptional level. However, given the proposed function of SINHCAF within the SIN3A repressor complex, we determined if the increases in HIF 2α protein we observed following SINHCAF depletion were observed at the mRNA level. Here, we analysed levels of HIF 1α and HIF 2α mRNA by qPCR in normoxia following SINHCAF depletion (Figure 3A). This analysis indicated that loss of SINHCAF resulted in a significant increase in HIF 2α RNA, but not HIF 1α (Figure 3A; Sup. Figure S2A). Similar results were also observed when SIN3A was depleted (Figure 3A). Given that the SIN3A is part of a repressor complex containing HDACs, we also extended our analysis to HDAC1. Here, we could confirm that both HIF 1α and HIF 2α mRNA levels were increased upon depletion of HDAC1 (Figure 3B), an effect that was also observed at the protein level (Figure 3B). Similar results were also obtained when cells were treated with the class I and II HDAC inhibitor trichostatin A (TSA) (Sup. Figure S2B-C). These results suggest that SIN3A/SINHCAF provide specificity in targeting HIF α isoforms.

**Figure 3:**
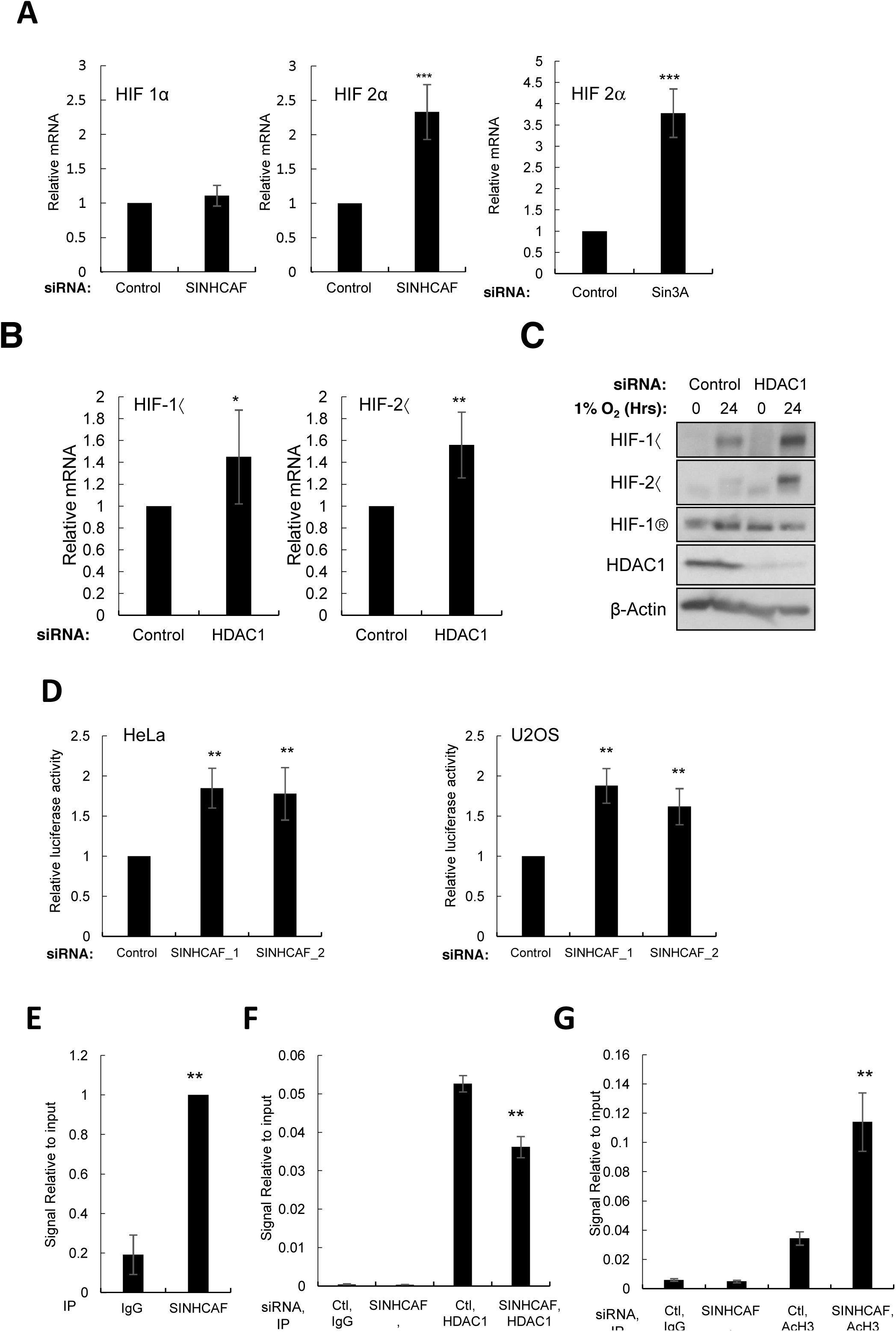
SINHCAF, but not HDAC1, is a specific repressor of HIF 2α promoter. (A) HeLa cells were transfected with Control, SINHCAF or SIN3A siRNA oligonucleotides for 48 hours prior to RNA extraction. RNA expression of the HIF-α isoforms was determined by qPCR. (B) HDAC1 or non-targeting siRNA oligonucleotides were transfected to HeLa cells prior to RNA extraction. RNA expression of the HIF-α isoforms was determined by qPCR. HDAC1 or non-targeting siRNA oligonucleotides were transfected to HeLa cells cultured in the presence or absence of hypoxia for 24 hours. Lysed samples were analysed by immunoblot for expression of HIF system isoforms and HDAC1. Graphs depict mean + S.E.M. * P ≤ 0.05, ** P ≤ 0.01, *** P ≤ 0.001. (D) HeLa and U2OS cells stably expressing an HIF 2α promoter-renilla luciferase reporter construct, were transfected with control or one of two SINHCAF [1/2] siRNA oligonucleotides and luciferase activity was measured. Graphs depict mean + S.E.M. * P ≤ 0.05, ** P ≤ 0.01, *** P ≤ 0.001. (E) ChIP for SINHCAF were performed in HeLa cells and HIF 2α promoter occupancy was analysed by qPCR. (F) ChIP for HDAC1 were performed in HeLa cells that had been transfected with Control or SINHCAF siRNA oligonucleotides. An assessment of HIF 2α promoter occupancy was performed by qPCR. (G) Change in histone H3 acetylation at the HIF 2α promoter was analysed following SINHCAF knockdown by qPCR. N=3. Graphs depict mean + S.E.M. * P ≤ 0.05, ** P ≤ 0.01, *** P ≤ 0.001. See also Figure S2.

To determine if the HIF 2α promoter was being specifically regulated by SINHCAF/SIN3A/ HDAC1 complex, we analysed HIF 2α promoter constructs in the presence or absence of SINHCAF. HeLa and U2OS cells, stably expressing a renilla luciferase-HIF 2α romoter construct, were transfected with control or one of two SINHCAF siRNA oligonucleotides (Figure 3D). In accordance with our previous results investigating HIF 2α mRNA levels, SINHCAF knockdown resulted in a significant increase in renilla luciferase activity, suggesting that the HIF 2α promoter is regulated by SINHCAF. A decrease in promoter activity was also observed when SINHCAF was overexpressed (Sup. Figure S2D).

To test if SINHCAF can control the HIF 2α promoter directly, as predicted by the ChIP-sequencing results, ChIP for SINHCAF and qPCR analysis using primers directed towards the HIF 2α promoter was performed. This analysis revealed a significant enrichment of SINHCAF present at the HIF 2α promoter (Figure 3E). To investigate if SINHCAF is important for the recruitment of HDAC1 to the HIF 2α promoter, and the functional consequences of this, we determined the levels of HDAC1 and AcH3 in the presence or absence of SINHCAF (Figure 3F-G). This analysis revealed that SINHCAF depletion resulted in a decreased in the levels of HDAC1 detected at the HIF 2α promoter (Figure 3F) with a correspondent increase in histone acetylation (Figure 3G) consistent with the proposed mechanism of SIN3A/HDAC1 recruitment to the HIF 2α promoter. Taken together, these results demonstrate that SINHCAF is important for HDAC1 recruitment to the HIF 2α promoter, controlling transcription of the HIF 2α gene.

### SINHCAF controls HIF 2α activity in cells

HIF transcription factors have important functions in tumour progression, such as controlling proliferation and angiogenesis (Ceradini et al., 2004; Gordan et al., 2007; Koshiji et al., 2004; Maltepe et al., 1997; Semenza, 2009). To determine the importance of SINHCAF mediated control of HIF 2α functions in cancer, in vitro cellular assays analysis of angiogenesis were performed. Human umbilical vein endothelial cell (HUVEC) tube formation is analogous to angiogenesis *in-vivo*. Conditioned media from DLD-1 tumour cells that had been treated with single or double siRNA knockdown of control, SINHCAF and HIF 2α and treated with hypoxia for 24 hours were collected. HUVECs were encouraged to form tubes in the presence of conditioned media for 24 hours. Loss of SINHCAF resulted in a significant increase in total tube length following treatment with conditioned media, loss of HIF 2α resulted in a significant reduction of the same parameter, and combined depletion recovered the effects observed with individual SINHCAF and HIF 2α depletions (Figure 4A).

**Figure 4:**
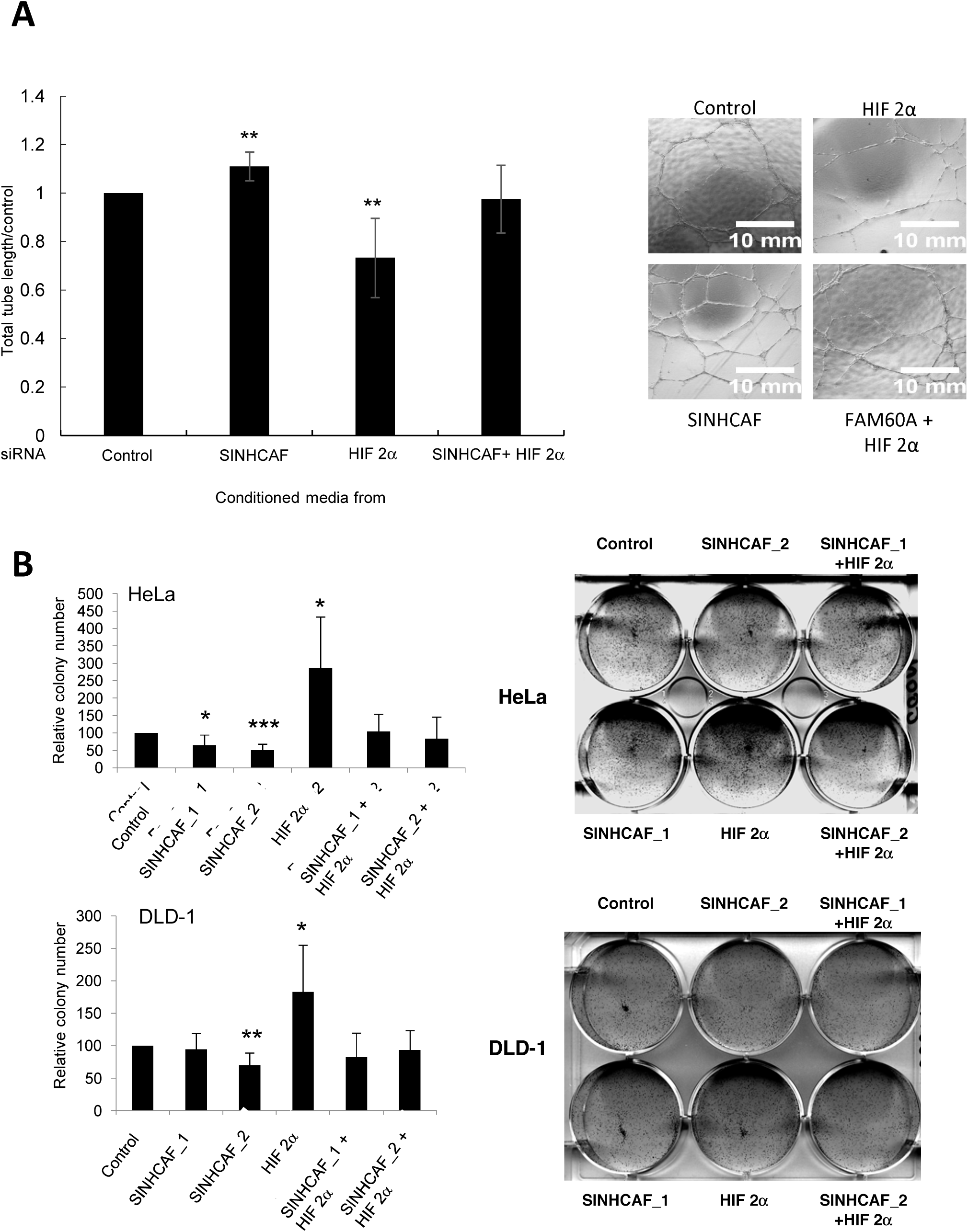
Functional Significance of SINHCAF-Mediated HIF 2α Repression. (A) Tube Formation: Conditioned media was collected from DLD-1 cells treated with single or double knockdown of control, SINHCAF or HIF 2α, and cultured in hypoxia for 24 hours. HUVEC were cultured in the presence of recombinant basement membrane and conditioned media for 24 hours. Total tube length was measured by Image J macros, representative images are shown. N=3. Graphs depict mean + S.E.M. * P ≤ 0.05, ** P ≤ 0.01, *** P ≤ 0.001. (B) Colony Formation: HeLa and DLD-1 cells were seeded for colony formation following siRNA transfection. Representative images are displayed. Relative colony number at day 7 for each cell line is shown. N-3. Graphs depict mean + S.E.M. * P ≤ 0.05, ** P ≤ 0.01, *** P ≤ 0.001.

An important cellular function deregulated in cancer is proliferation. To determine how SINHCAF control over HIF 2α alters cell proliferation, HeLa and DLD-1 cells were transfected with control, SINHCAF, HIF 2α, or SINHCAF and HIF 2α siRNA’s, and cell proliferation was analysed using colony-forming assays. This analysis revealed that SINHCAF depletion resulted in decreased number of colonies, while HIF 2α depletion significantly increased colony formation in both cell types (Figure 4B). Importantly, the effect of SINHCAF depletion on colony numbers was shown to be dependent on HIF 2α (Figure 4B), demonstrating the importance of SINHCAF control over HIF 2α levels and function.

### SINHCAF mRNA levels are elevated in human cancer and they are inversely correlated with HIF 2α

Analysis of the publically available datasets on Oncomine found SINHCAF to be frequently upregulated in colorectal cancer but also other cancer types such as Leukemia and Lung cancer (Figure 5A), interestingly, levels of HIF 2α mRNA are often reduced in a variety of cancer such as colorectal and lung (Figure 5A). To understand if there is any biological significance to the elevated level of SINHCAF and decreased level of HIF 2α mRNA, we analysed publicly available patient datasets for correlation of the levels of these genes and patient survival (Figure 5B-C). We used KM plot (Gyorffy et al., 2013) for lung cancer dataset and prognoscan for colorectal cancer (Mizuno et al., 2009). This analysis revealed that high levels of SINHCAF correlated with poor patient survival with a Hazzard Ratio of 1.78 in lung and 2.14 in colorectal cancer. On the other hand, high levels of HIF 2α correlated with increased patient survival in both types of cancer (Figure 5B-C).

**Figure 5:**
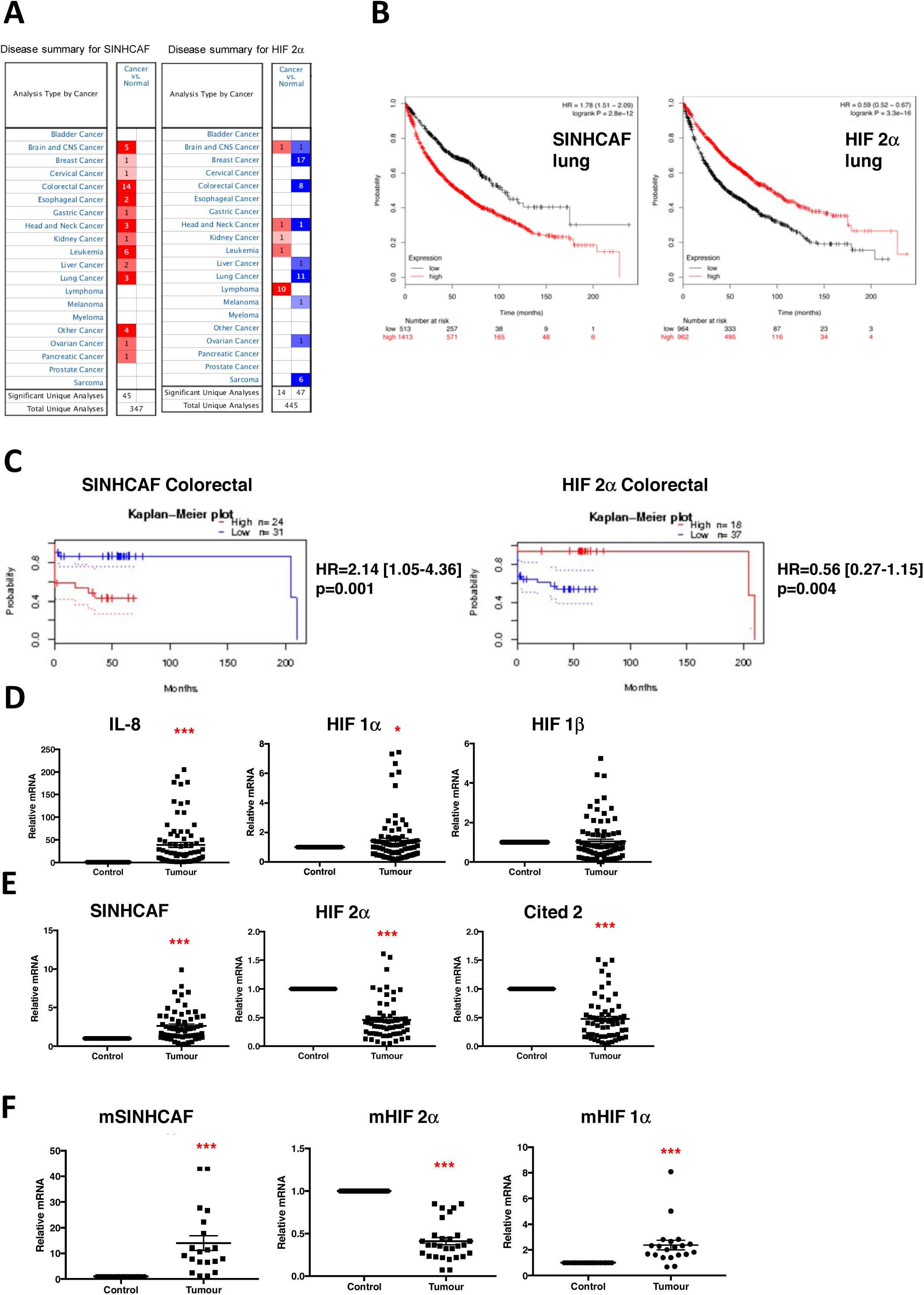
SINHCAF and HIF 2α levels are inversely correlated in cancer and have converse prognostic value. (A) Cancer summary for mRNA levels of SINHCAF and HIF 2a across different cancer types based on publish datasets in Oncomine. (B) Kaplan-Mayer plot for SINHCAF and HIF 2α mRNA in lung cancer HR-Hazzard Ratio. (C) Kaplan-Mayer plot for SINHCAF and HIF 2α mRNA in colorectal cancer HR-Hazzard Ratio. (D) RNA was extracted from fresh frozen tissue taken from 45 patients suffering from colorectal cancer of known stage and expression of IL-8, HIF-1α, and HIF-1β was determined by qPCR. (E) as in D, but levels of SINHCAF, HIF 2α and Cited 2 mRNA were determined by qPCR. Paired samples of healthy tissue from each patient were used as control. Scatter = Mean +/-S.E.M. * P ≤ 0.05, ** P ≤ 0.01, *** P ≤ 0.001. F. RNA was extracted from fresh frozen tissue taken from 10 individual Apc(Min/+) Gstp1/p2(-/-) mice, levels of SINHCAF, HIF 1α, and HIF 2β were determined by qPCR. Scatter = Mean +/-S.E.M. * P ≤ 0.05, ** P ≤ 0.01, *** P ≤ 0.001.

To validate our bioinformatics analysis, fresh samples of 45 colorectal cancers with paired normal controls were procured from the Tayside Tissue Bank. Stratification of colorectal cancer according to the Duke’s staging system was performed and RNA was extracted from a total of 90 samples (45 cancer / 45 normal). The patient demographics are summarised in Table 1. RNA extracted from these cancers was examined by qPCR for expression of the inflammatory cytokine IL-8 (Figure 5D), as its expression can be predictive of colorectal cancer stage (Rubie et al., 2007. A significant increase in IL-8 was observed when tumour tissue was compared to normal match control tissue. Interestingly, a significant increase in HIF 1α mRNA but no significant different in HIF 1β mRNA was observed in these patients (Figure 5D). HIF 1α RNA can be induced at late stages of the disease {Yoshimura, 2004 #212). Importantly expression of SINHCAF mRNA was found to be significantly higher in this patient cohort with a correspondent reduced level of HIF 2α and its target Cited-2 (Figure 5E).

We extended our analysis of SINHCAF and HIF 2α regulation to a mouse model of colon cancer (Ritchie et al., 2009). Here, APC (Min/+) mice with a simultaneous deletion of glutathione transferase Pi Gstp1/p2(-/-), develop colorectal tumours which are characterised by increased inflammation, closely mimicking the human colon cancer situation {Ritchie). Tumours and normal matched control tissue from the colon of these mice were analysed for the mRNA levels of HIF 2α, HIF 1α and SINHCAF (Figure 5F). As observed using the human patient material, levels of HIF 1α were higher in the tumour than in normal tissue. Conversely, and as predicted from analysis of the human data, HIF 2α levels were reduced while levels of SINHCAF were significantly elevated in the mouse tumour samples.

Taken together, these results indicate that SINHCAF is a putative biomarker for colorectal and lung cancer. Furthermore, our data indicates that control of HIF 2α is an important factor determining cancer progression in these types of tumours.

## Discussion

In this study, we have used genome-wide approaches and an integrative analysis of publicly available datasets to identify the biological functions of SINHCAF, a relatively unknown protein. We have found that SINHCAF function correlates with a hypoxic and inflammatory signature in cancer cells. We further describe a novel mechanism controlling HIF 2α levels and function with important implications for human cancer. We found that SINHCAF acts to epigenetically silence expression of the HIF 2α gene through recruitment of the class I SIN3-HDAC co-repressor factor. SINHCAF expression is linked to functional cellular changes in angiogenesis (tube formation), and proliferation *in-vitro*. Importantly, SINHCAF and HIF 2α levels inversely correlate with patient survival in both lung and colorectal cancer. Finally, this inverse correlation in expression is also observed in a genetic mouse model for colon cancer.

Our study has also revealed that SIN3A occupancy across the genome is cell type specific, with only a subset of genes are occupied by SIN3A (which we called core/conserved SIN3A regulated genes) in all the cell backgrounds we analysed. While preparing this manuscript, an interesting study was published concerning the function of SINHCAF in mouse stem cells. There, SINHCAF was shown to be required for maintaining SIN3A function in a subset of genes, and preventing unnecessary differentiation {Streubel, 2017}. This study has suggested that SINHCAF and SIN3A incorporate at 1:1 stoichiometry only in ES stems, but not in differentiated cells. Our data are in agreement with this possibility, as our ChIP-seq analysis revealed that only the core function of SIN3A across cell types is shared with SINHCAF at a rate of 94% overlap, while cell type differences in gene expression vary from 28-60% overlap between these two factors.

Evidence is emerging to suggest that control of the basal expression of the HIF system is as important as its hypoxia-responsive post-translational degradation (Kuschel et al., 2012). Changes in basal expression of the HIF α isoforms are important in determining tissue-specific, hypoxia-inducible gene expression and are also important in the progression of multiple types of disease including cancer (Bandarra et al., 2014; Blouin et al., 2004; Gorlach, 2009; Kenneth et al., 2009; Kuschel et al., 2012; Page et al., 2002; van Uden et al., 2008; van Uden et al., 2011). Basal HIF expression appears to be regulated by a transcriptional rheostat that is designed to allow an organism to respond in a heterogeneous manner to changes in oxygen supply and demand through space and time. The HIF system is set differently in different cells, in a manner that is appropriate for the physiological control of oxygen homeostasis (Rosenberger et al., 2002). Our own laboratory has demonstrated the importance of regulation of the expression of HIF 1α previously. We demonstrated that the ATP-dependent chromatin remodeling complex SWI/SNF directly affects both the expression of HIF 1α, and its ability to transactivate target genes (Kenneth et al., 2009).

The specific nature of the mechanism involving SINHCAF would allow selective repression of HIF 2α. This proposed mechanism of transcriptional regulation by chromatin modification is one established mechanism for the heritable control of transcription that would be capable of specifically controlling the temporal and spatial transcription of HIF 2α, a phenomenon already seen in several cell types. SINHCAF repression was shown to be specific for HIF 2α, while HDAC1 repression is seen for both HIF 2α and HIF 1α. SINHCAF thus acts as a specificity factor for HDAC-mediated control of the HIF genes.

SINHCAF importance is exemplified by the level of deregulation in human cancer. Analysis of publicly available datasets present on oncomine, demonstrated that in colorectal cancer, over 40% of datasets have overexpressed SINHCAF. In addition, cancers such as lung, also have a significant level of SINHCAF overexpression. Furthermore, SINHCAF were shown to be predictive of patient survival for both colon and lung cancer, further demonstrating the importance of this protein. Our own analysis, demonstrated a correlation between levels of SINHCAF and colorectal cancer disease appearance. HIF 2α repression has also been shown to be important for progression of the disease (Rawluszko-Wieczorek et al., 2014), and Cited-2 loss has been shown to increase colon cancer cell invasiveness through induction of MMP-13 in an HDAC-dependent mechanism (Bai and Merchant, 2007). This reciprocal relationship between SINHCAF and HIF 2α was maintained in patient samples we analysed, and both SINHCAF, HIF 2α and the HIF 2α-target gene Cited-2 were identified as biomarkers with potential use for the early diagnosis of colorectal cancer.

*In-vitro* analysis of SINHCAF and HIF 2α regulated cellular responses demonstrated their co-dependency. Our results demonstrate that indeed, SINHCAF depletion prevents colony formation, while HIF 2α depletion induced a significant increase of colonies formed. Again, these responses were reversed upon co-depletion. Furthermore, SINHCAF was seen to repress tube formation via HIF 2α with total recovery achieved following co-depletion. This implies that SINHCAF depletion results in an increased quality of blood vessels, an important aspect required for tumour suppression (Semenza, 2013). HIF 2α repression can drive tumour progression indirectly through loss of expression of HIF 2α dependent tumour suppressors. In support of these proposals, HIF 2α but not HIF 1α deletion has been shown to increase tumour growth and progression secondary to loss of the tumor suppressor secretoglobin 3a1 protein in a KRAS driven lung tumour model (Krop et al., 2005; Mazumdar et al., 2010).

In colorectal cancer, we observed increased expression of HIF 1α in keeping with work by others that has identified HIF 1α expression as prognostically poor in terms of disease outcome (Baba et al., 2010; Saigusa et al., 2012; Yoshimura et al., 2004). Given the increase observed in IL-8, it is possible that HIF 1α RNA increases via a NF-κB dependent mechanism (van Uden et al., 2008). However, further work is necessary to investigate these possibilities.

The mechanism for the epigenetic regulation of HIF 2α by SINHCAF described in this manuscript represents an exciting and potentially druggable future therapeutic target for the treatment of colorectal cancer. In the physiological environment, these mechanisms may play a more important role in the tissue specific expression and kinetic response of the HIF α isoforms than is currently understood. Selective inhibition of a specific HIF α isoform in the context of cancer is an attractive proposal because it would allow enhanced specificity and targeting of therapy. Appropriate focus on the development of small molecule inhibitors of SINHCAF would provide an important therapy to reverse the effects of this protein whose deregulation is implicated in multiple disease processes. It might also be useful in predicting which patients might respond to HDAC inhibitors, which are already developed but have received conflicting results in clinical trials (Nervi et al., 2015).

Importantly, these findings are not restricted in application to a cancer model. We have demonstrated SINHCAF as an important protein in the control of pathological responses associated with hypoxia and inflammation, and shown that SINHCAF is a specific epigenetic regulator of HIF 2α. These features make the mechanism described relevant to multiple physiological and pathological processes that involve hypoxia and inflammation and therefore have wide importance across a range of human diseases.

## Methods

### Cell Culture and Transfection

All cell lines were maintained for no more than 30 passages and grown in Dulbecco’s modified Eagle’s medium or Roswell Park Memorial Institute medium containing 1% penicillin and streptomycin, supplemented with 10% fetal bovine serum (FBS). Hypoxic treatments were 1% O2 unless otherwise stated and delivered using a Baker-Ruskinn InVivo2 300 hypoxia chamber. All expression plasmids and siRNA oligonucleotides are shown in the Supplemental Experimental Procedures. siRNA transfections were performed using Interferin (Peqlab), and DNA transfections using TurboFect (Thermo). All reagents were used according to manufacturer’s instructions.

### SINHCAF ChIP-sequencing and analysis

DNA from each ChIP sample and corresponding input (three replicates per condition) were used in paired end DNA library construction and sequenced on a HiSeq 2000 platform (Illumina) at the University of Dundee Tayside Centre for Genomic Analysis. Fastq data files were passed through the quality control tool FASTQC and aligned to the human genome annotation hg19_73 using Bowtie 2 (Langmead et al., 2009). The data was then converted to bam (individual replicates and merged) and subsequently converted to BED and bigwig file formats containing mapped and unmapped read count information using R Bioconductor packages GenomicRanges (Lawrence et al., 2013), Repitools (Statham et al., 2010) and rtracklayer (Lawrence et al., 2009). Peaks for individual replicates and merged bam files were called using the MACS2 software in narrow peak mode with input bam files as controls and a FDR cut off of 0.005. Peaks called in only 1 out of 3 replicates were discarded from analysis. R Bioconductor packages were used for overlapping peaks (ChIPpeakAnno) (Zhu et al., 2010), identification of the nearest gene for each peak (bioMart) (Durinck et al., 2009), (ChIPseeker) (Yu et al., 2015)), peak genomic annotation assignment (ChIPseeker), generation of library size normalised peak read counts vs transcription start site figures (CoverageView) and peak coverage track images (Gvis) (Hahne and Ivanek, 2016). For genomic annotation assignment, a promoter is defined as between 3kb upstream and downstream of a TSS and an intergenic region is defined as greater than 3kb downstream of the nearest TES and greater than 3kb upstream of the nearest TSS. The following ChIP sequencing datasets from the encode project (Consortium, 2012; Sloan et al., 2016) were downloaded from the NCBI GEO database, Hela S3 H3K9ac (GSM733756), Hela S3 Pol2 (GSM733759), hESC SIN3A (GSM935289), MCF7 SIN3A (GSM1010862), A549 SIN3A (GSM1010882), HCT116 SIN3A (GSM1010905).

### Integrative analysis using public datasets

Analysis of A549 microarray (Smith et al., 2012) was performed using GEO2R tool on the GEO website. Analysis of hallmark gene signatures was performed using the MSigDB from Gene Set Enrichment Analysis software (Mootha et al., 2003; Subramanian et al., 2005). Analysis of lung patient datasets for the prognostic value of SINHCAF and HIF 2α was done using KM plot tool (Gyorffy et al., 2013). Prognostic value for SINHCAF and HIF 2α in colorectal cancer patient datasets was performed using the Prognoscan database (Mizuno et al., 2009) based on the published data (Freeman et al., 2012; Smith et al., 2010). In addition, levels of SINHCAF and HIF 2α in cancer were obtained analyzing the Oncomine database.

### qRT-PCR, siRNA, Luciferase assay, Western Blots and Immunoprecipitations

Additional experimental procedures, such as immunoprecipitations and luciferase assays, have been described previously (Kenneth et al., 2013).

### ChIP-PCR assays

ChIP assays were performed as described previously (Schumm et al., 2006).

### Tube Formation Assay

Tube formation assays were performed using HUVECs and μ-slides (i-Bidi) as described in (Bandarra et al., 2015).

### Colony Formation Assay

HeLa and DLD-1 cells were transfected with control or SINHCAF, HIF 2α or a combination of the two siRNAs for 24 h prior to trypsination. Cells were counted and plates on two 6 well plates at the density of 5000 or 10000 cells per well. Cells were allowed to grow for 6 days prior to staining with Crystal Violet. Plates were imaged and colonies counted using Image J.

### Patient Cohort

We obtained fresh frozen tumour specimens of colorectal cancer patients, which were histopathologically diagnosed in the Department of Pathology, Ninewells Hospital, Dundee, Scotland. All tumours were primary and were untreated before surgery. The corresponding survival status was confirmed by means of patient or family contact. The overall survival duration was defined from the date of surgical resection to death or last known date alive. After giving written consent, demographic and clinical data were collected in the hospitals using a standard interviewer administered questionnaire and/or medical records. Surgery-removed samples of all cases were collected. This study was approved by the Tayside tissue bank, Ninewells Hospital, Dundee, Scotland. Informed consent was obtained from all subjects or their relatives.

### Animals

Experiments were undertaken in accordance with the Animals (Scientific Procedures) Act of 1986. The animal study plan was developed after ethical approval was granted (Project Licence 60/5986), and was further approved by the Named Veterinary Surgeon and the Director of Biological Services of the University of Dundee. Apc(Min/+) Gstp1/p2(-/-) mice on a C56BL/6 background (initially obtained from Dr Colin Henderson and Professor C. Roland Wolf, University of Dundee) were bred and maintained in the Medical School Resource Unit of the University of Dundee on a 12-h light/ 12-h dark cycle and 35% humidity. Throughout the study, the animals had free access to water and pelleted RM1 diet (SDS Ltd., Witham, Essex, UK). The animals were monitored regularly for symptoms of intestinal neoplasia and general ill health throughout the study.

### Mouse tissue and RNA extraction

Male Apc(Min/+) Gstp1/p2(-/-) mice [*Gstp*-null *Apc*^*Min*^] were sacrificed at 20 weeks of age by CO_2_ asphyxiation. Colon tissue was dissected on ice and flushed with ice-cold PBS. Colon tumours and surrounding normal tissue were excised and flash-frozen in liquid N_2_ followed by storage at -80 °C. Tumour and matched normal samples were processed for RNA extraction using RNAesy Kit from Qiagen.

### Statistical Analysis

Data analysis was performed using SigmaPlot v12.0 (Systat Software Inc., California, U.S.A.). One-way analysis of variance (ANOVA) with Holm-Sidak pairwise comparison was used to compare results between groups, data are presented as category mean +/-standard error of the mean unless otherwise stated. All statistical tests were two sided and statistical significance denoted as follows: * p ≤ 0.05, ** p ≤ 0.01, *** p ≤ 0.001.

## Author Contributions

J.R. and S.R. initiated the project and analysed the data; J.B., M.B., A.S., D.B., I.M., R.S. E.K. and S.R. performed experiments and additional biochemical analysis; and J.B., T.O.H., A.D.K. and S.R. wrote the manuscript.

## Acknowledgements

We would like to thank Prof. John Rouse for providing reagents and critical reading of the manuscript. J.B was funded by a Cancer Research UK clinical fellowship; D.B. was funded by a PhD studentship from the Portuguese Science Foundation and Graduate Program in Areas of Basic and Applied Biology (GABBA); M.B. is funded by a MRC PhD studentship. A.S. was funded by a Wellcome Trust PhD studentship. S.R. is funded by a Cancer Research UK Senior Research Fellowship C99667/A12918; TOH is funded by a Wellcome Trust Senior Fellowship to TOH (095062). E.K. and A.D.K are supported by Cancer Research UK (C20953/A18644). This work was also supported by a Wellcome Trust Strategic Award (097945/B/11/Z) and additionally supported by a small grant from the Anne and Normile Baxter prize for cancer research 2014.

## Supplementary Information accompanies the paper

## References

Baba, Y., Nosho, K., Shima, K., Irahara, N., Chan, A.T., Meyerhardt, J.A., Chung, D.C., Giovannucci, E.L., Fuchs, C.S., and Ogino, S. (2010). HIF1A overexpression is associated with poor prognosis in a cohort of 731 colorectal cancers. Am J Pathol 176, 2292–2301.

Bai, L., and Merchant, J.L. (2007). A role for CITED2, a CBP/p300 interacting protein, in colon cancer cell invasion. FEBS Lett 581, 5904–5910.

Bandarra, D., Biddlestone, J., Mudie, S., Muller, H.A., and Rocha, S. (2014). Hypoxia activates IKK-NF-kappaB and the immune response in Drosophila melanogaster. Biosci Rep 34.

Bandarra, D., Biddlestone, J., Mudie, S., Muller, H.A., and Rocha, S. (2015). HIF-1alpha restricts NF-kappaB-dependent gene expression to control innate immunity signals. Dis Model Mech 8, 169–181.

Belaiba, R.S., Bonello, S., Zahringer, C., Schmidt, S., Hess, J., Kietzmann, T., and Gorlach, A. (2007). Hypoxia up-regulates hypoxia-inducible factor-1alpha transcription by involving phosphatidylinositol 3-kinase and nuclear factor kappaB in pulmonary artery smooth muscle cells. Mol Biol Cell 18, 4691–4697.

Biddlestone, J., Bandarra, D., and Rocha, S. (2015). The role of hypoxia in inflammatory disease (review). International journal of molecular medicine 35, 859–869.

Biswas, S., Mukherjee, R., Tapryal, N., Singh, A.K., and Mukhopadhyay, C.K. (2013). Insulin regulates hypoxia-inducible factor-1alpha transcription by reactive oxygen species sensitive activation of Sp1 in 3T3-L1 preadipocyte. PLoS One 8, e62128.

Blouin, C.C., Page, E.L., Soucy, G.M., and Richard, D.E. (2004). Hypoxic gene activation by lipopolysaccharide in macrophages: implication of hypoxia-inducible factor 1alpha. Blood 103, 1124–1130.

Bonello, S., Zahringer, C., BelAiba, R.S., Djordjevic, T., Hess, J., Michiels, C., Kietzmann, T., and Gorlach, A. (2007). Reactive oxygen species activate the HIF-1alpha promoter via a functional NFkappaB site. Arterioscler Thromb Vasc Biol 27, 755–761.

Bruegge, K., Jelkmann, W., and Metzen, E. (2007). Hydroxylation of hypoxia-inducible transcription factors and chemical compounds targeting the HIF-alpha hydroxylases. Curr Med Chem 14, 1853–1862.

Carroll, V.A., and Ashcroft, M. (2006). Role of hypoxia-inducible factor (HIF)-1alpha versus HIF-2alpha in the regulation of HIF target genes in response to hypoxia, insulin-like growth factor-I, or loss of von Hippel-Lindau function: implications for targeting the HIF pathway. Cancer Res 66, 6264–6270.

Ceradini, D.J., Kulkarni, A.R., Callaghan, M.J., Tepper, O.M., Bastidas, N., Kleinman, M.E., Capla, J.M., Galiano, R.D., Levine, J.P., and Gurtner, G.C. (2004). Progenitor cell trafficking is regulated by hypoxic gradients through HIF-1 induction of SDF-1. Nat Med 10, 858–864.

Consortium, E.P. (2012). An integrated encyclopedia of DNA elements in the human genome. Nature 489, 57–74.

Cruzeiro, G.A., Dos Reis, M.B., Silveira, V.S., Lira, R.C., Carlotti, C.G., Neder, L., Oliveira, R.S., Yunes, J.A., Brandalise, S.R., Aguiar, S., et al. (2017). HIF1A is overexpressed in medulloblastoma and its inhibition reduces proliferation and increases EPAS1 and ATG16L1 methylation. Curr Cancer Drug Targets.

Cui, J., Duan, B., Zhao, X., Chen, Y., Sun, S., Deng, W., Zhang, Y., Du, J., Chen, Y., and Gu, L. (2016). MBD3 mediates epigenetic regulation on EPAS1 promoter in cancer. Tumour Biol 37, 13455–13467.

D’Ignazio, L., Bandarra, D., and Rocha, S. (2016). NF-kappaB and HIF crosstalk in immune responses. FEBS J 283, 413–424.

D’Ignazio, L., Batie, M., and Rocha, S. (2017). Hypoxia and Inflammation in Cancer, Focus on HIF and NF-kappaB. Biomedicines 5.

Dong, G., Mao, Q., Yu, D., Zhang, Y., Qiu, M., Dong, G., Chen, Q., Xia, W., Wang, J., Xu, L., et al. (2017). Integrative analysis of copy number and transcriptional expression profiles in esophageal cancer to identify a novel driver gene for therapy. Sci Rep 7, 42060.

Durinck, S., Spellman, P.T., Birney, E., and Huber, W. (2009). Mapping identifiers for the integration of genomic datasets with the R/Bioconductor package biomaRt. Nat Protoc 4, 1184–1191.

Epstein, A.C., Gleadle, J.M., McNeill, L.A., Hewitson, K.S., O’Rourke, J., Mole, D.R., Mukherji, M., Metzen, E., Wilson, M.I., Dhanda, A., et al. (2001). C. elegans EGL-9 and mammalian homologs define a family of dioxygenases that regulate HIF by prolyl hydroxylation. Cell 107, 43–54.

Fandrey, J., Gorr, T.A., and Gassmann, M. (2006). Regulating cellular oxygen sensing by hydroxylation. Cardiovasc Res 71, 642–651.

Frede, S., Stockmann, C., Freitag, P., and Fandrey, J. (2006). Bacterial lipopolysaccharide induces HIF-1 activation in human monocytes via p44/42 MAPK and NF-kappaB. Biochem J 396, 517–527.

Freeman, T.J., Smith, J.J., Chen, X., Washington, M.K., Roland, J.T., Means, A.L., Eschrich, S.A., Yeatman, T.J., Deane, N.G., and Beauchamp, R.D. (2012). Smad4-mediated signaling inhibits intestinal neoplasia by inhibiting expression of beta-catenin. Gastroenterology 142, 562–571 e562.

Gordan, J.D., Thompson, C.B., and Simon, M.C. (2007). HIF and c-Myc: sibling rivals for control of cancer cell metabolism and proliferation. Cancer Cell 12, 108–113.

Gorlach, A. (2009). Regulation of HIF-1alpha at the transcriptional level. Curr Pharm Des 15, 3844–3852.

Gyorffy, B., Surowiak, P., Budczies, J., and Lanczky, A. (2013). Online survival analysis software to assess the prognostic value of biomarkers using transcriptomic data in non-small-cell lung cancer. PLoS One 8, e82241.

Hahne, F., and Ivanek, R. (2016). Visualizing Genomic Data Using Gviz and Bioconductor. Methods Mol Biol 1418, 335–351.

Hu, C.J., Iyer, S., Sataur, A., Covello, K.L., Chodosh, L.A., and Simon, M.C. (2006). Differential regulation of the transcriptional activities of hypoxia-inducible factor 1 alpha (HIF-1alpha) and HIF-2alpha in stem cells. Mol Cell Biol 26, 3514–3526.

Imamura, M., Takahashi, A., Yamauchi, T., Hara, K., Yasuda, K., Grarup, N., Zhao, W., Wang, X., Huerta-Chagoya, A., Hu, C., et al. (2016). Genome-wide association studies in the Japanese population identify seven novel loci for type 2 diabetes. Nature communications 7, 10531.

Ivan, M., Kondo, K., Yang, H., Kim, W., Valiando, J., Ohh, M., Salic, A., Asara, J.M., Lane, W.S., and Kaelin, W.G., Jr. (2001). HIFalpha targeted for VHL-mediated destruction by proline hydroxylation: implications for O2 sensing. Science 292, 464–468.

Jaakkola, P., Mole, D.R., Tian, Y.M., Wilson, M.I., Gielbert, J., Gaskell, S.J., von Kriegsheim, A., Hebestreit, H.F., Mukherji, M., Schofield, C.J., et al. (2001). Targeting of HIF-alpha to the von Hippel-Lindau ubiquitylation complex by O2-regulated prolyl hydroxylation. Science 292, 468–472.

Kadamb, R., Mittal, S., Bansal, N., Batra, H., and Saluja, D. (2013). Sin3: insight into its transcription regulatory functions. Eur J Cell Biol 92, 237–246.

Kaelin, W.G., Jr., and Ratcliffe, P.J. (2008). Oxygen sensing by metazoans: the central role of the HIF hydroxylase pathway. Mol Cell 30, 393–402.

Kazi, A.A., Gilani, R.A., Schech, A.J., Chumsri, S., Sabnis, G., Shah, P., Goloubeva, O., Kronsberg, S., and Brodie, A.H. (2014). Nonhypoxic regulation and role of hypoxia-inducible factor 1 in aromatase inhibitor resistant breast cancer. Breast Cancer Res 16, R15.

Keith, B., and Simon, M.C. (2007). Hypoxia-inducible factors, stem cells, and cancer. Cell 129, 465–472.

Kenneth, N.S., Mudie, S., Naron, S., and Rocha, S. (2013). TfR1 interacts with the IKK complex and is involved in IKK-NF-kappaB signalling. Biochem J 449, 275–284.

Kenneth, N.S., Mudie, S., van Uden, P., and Rocha, S. (2009). SWI/SNF regulates the cellular response to hypoxia. J Biol Chem 284, 4123–4131.

Koshiji, M., Kageyama, Y., Pete, E.A., Horikawa, I., Barrett, J.C., and Huang, L.E. (2004). HIF-1alpha induces cell cycle arrest by functionally counteracting Myc. EMBO J 23, 1949–1956.

Koshikawa, N., Hayashi, J., Nakagawara, A., and Takenaga, K. (2009). Reactive oxygen species-generating mitochondrial DNA mutation up-regulates hypoxia-inducible factor-1alpha gene transcription via phosphatidylinositol 3-kinase-Akt/protein kinase C/histone deacetylase pathway. J Biol Chem 284, 33185–33194.

Krop, I., Parker, M.T., Bloushtain-Qimron, N., Porter, D., Gelman, R., Sasaki, H., Maurer, M., Terry, M.B., Parsons, R., and Polyak, K. (2005). HIN-1, an inhibitor of cell growth, invasion, and AKT activation. Cancer Res 65, 9659–9669.

Kuschel, A., Simon, P., and Tug, S. (2012). Functional regulation of HIF-1alpha under normoxia--is there more than post-translational regulation? J Cell Physiol 227, 514–524.

Lafleur, V.N., Richard, S., and Richard, D.E. (2014). Transcriptional repression of hypoxia-inducible factor-1 (HIF-1) by the protein arginine methyltransferase PRMT1. Mol Biol Cell 25, 925–935.

Langmead, B., Trapnell, C., Pop, M., and Salzberg, S.L. (2009). Ultrafast and memory-efficient alignment of short DNA sequences to the human genome. Genome Biol 10, R25.

Lawrence, M., Gentleman, R., and Carey, V. (2009). rtracklayer: an R package for interfacing with genome browsers. Bioinformatics 25, 1841–1842.

Lawrence, M., Huber, W., Pages, H., Aboyoun, P., Carlson, M., Gentleman, R., Morgan, M.T., and Carey, V.J. (2013). Software for computing and annotating genomic ranges. PLoS Comput Biol 9, e1003118.

Maltepe, E., Schmidt, J.V., Baunoch, D., Bradfield, C.A., and Simon, M.C. (1997). Abnormal angiogenesis and responses to glucose and oxygen deprivation in mice lacking the protein ARNT. Nature 386, 403–407.

Mazumdar, J., Hickey, M.M., Pant, D.K., Durham, A.C., Sweet-Cordero, A., Vachani, A., Jacks, T., Chodosh, L.A., Kissil, J.L., Simon, M.C., et al. (2010). HIF-2alpha deletion promotes Kras-driven lung tumor development. Proc Natl Acad Sci U S A 107, 14182–14187.

Mizuno, H., Kitada, K., Nakai, K., and Sarai, A. (2009). PrognoScan: a new database for meta-analysis of the prognostic value of genes. BMC Med Genomics 2, 18.

Moniz, S., Bandarra, D., Biddlestone, J., Campbell, K.J., Komander, D., Bremm, A., and Rocha, S. (2015). Cezanne regulates E2F1-dependent HIF2alpha expression. J Cell Sci 128, 3082–3093.

Moniz, S., Biddlestone, J., and Rocha, S. (2014). Grow(2): the HIF system, energy homeostasis and the cell cycle. Histol Histopathol 29, 589–600.

Mootha, V.K., Lindgren, C.M., Eriksson, K.F., Subramanian, A., Sihag, S., Lehar, J., Puigserver, P., Carlsson, E., Ridderstrale, M., Laurila, E., et al. (2003). PGC-1alpha-responsive genes involved in oxidative phosphorylation are coordinately downregulated in human diabetes. Nature genetics 34, 267–273.

Munoz, I.M., MacArtney, T., Sanchez-Pulido, L., Ponting, C.P., Rocha, S., and Rouse, J. (2012). Family with sequence similarity 60A (FAM60A) protein is a cell cycle-fluctuating regulator of the SIN3-HDAC1 histone deacetylase complex. J Biol Chem 287, 32346–32353.

Nakazawa, M.S., Eisinger-Mathason, T.S., Sadri, N., Ochocki, J.D., Gade, T.P., Amin, R.K., and Simon, M.C. (2016). Epigenetic re-expression of HIF-2alpha suppresses soft tissue sarcoma growth. Nature communications 7, 10539.

Nervi, C., De Marinis, E., and Codacci-Pisanelli, G. (2015). Epigenetic treatment of solid tumours: a review of clinical trials. Clin Epigenetics 7, 127.

Niu, G., Briggs, J., Deng, J., Ma, Y., Lee, H., Kortylewski, M., Kujawski, M., Kay, H., Cress, W.D., Jove, R., et al. (2008). Signal transducer and activator of transcription 3 is required for hypoxia-inducible factor-1alpha RNA expression in both tumor cells and tumor-associated myeloid cells. Mol Cancer Res 6, 1099–1105.

Oh, Y.T., Lee, J.Y., Yoon, H., Lee, E.H., Baik, H.H., Kim, S.S., Ha, J., Yoon, K.S., Choe, W., and Kang, I. (2008). Lipopolysaccharide induces hypoxia-inducible factor-1 alpha mRNA expression and activation via NADPH oxidase and Sp1-dependent pathway in BV2 murine microglial cells. Neurosci Lett 431, 155–160.

Page, E.L., Robitaille, G.A., Pouyssegur, J., and Richard, D.E. (2002). Induction of hypoxia-inducible factor-1alpha by transcriptional and translational mechanisms. J Biol Chem 277, 48403–48409.

Patel, N., and Kalra, V.K. (2010). Placenta growth factor-induced early growth response 1 (Egr-1) regulates hypoxia-inducible factor-1alpha (HIF-1alpha) in endothelial cells. J Biol Chem 285, 20570–20579.

Patel, S.A., and Simon, M.C. (2008). Biology of hypoxia-inducible factor-2alpha in development and disease. Cell Death Differ 15, 628–634.

Pugh, C.W., Tan, C.C., Jones, R.W., and Ratcliffe, P.J. (1991). Functional analysis of an oxygen-regulated transcriptional enhancer lying 3’ to the mouse erythropoietin gene. Proc Natl Acad Sci U S A 88, 10553–10557.

Rawluszko-Wieczorek, A.A., Horbacka, K., Krokowicz, P., Misztal, M., and Jagodzinski, P.P. (2014). Prognostic Potential of DNA Methylation and Transcript Levels of HIF1A and EPAS1 in Colorectal Cancer. Mol Cancer Res.

Ritchie, K.J., Walsh, S., Sansom, O.J., Henderson, C.J., and Wolf, C.R. (2009). Markedly enhanced colon tumorigenesis in Apc(Min) mice lacking glutathione S-transferase Pi. Proc Natl Acad Sci U S A 106, 20859–20864.

Rius, J., Guma, M., Schachtrup, C., Akassoglou, K., Zinkernagel, A.S., Nizet, V., Johnson, R.S., Haddad, G.G., and Karin, M. (2008). NF-kappaB links innate immunity to the hypoxic response through transcriptional regulation of HIF-1alpha. Nature 453, 807–811.

Rocha, S. (2007). Gene regulation under low oxygen: holding your breath for transcription. Trends Biochem Sci 32, 389–397.

Rosenberger, C., Mandriota, S., Jurgensen, J.S., Wiesener, M.S., Horstrup, J.H., Frei, U., Ratcliffe, P.J., Maxwell, P.H., Bachmann, S., and Eckardt, K.U. (2002). Expression of hypoxia-inducible factor-1alpha and -2alpha in hypoxic and ischemic rat kidneys. J Am Soc Nephrol 13, 1721–1732.

Rubie, C., Frick, V.O., Pfeil, S., Wagner, M., Kollmar, O., Kopp, B., Graber, S., Rau, B.M., and Schilling, M.K. (2007). Correlation of IL-8 with induction, progression and metastatic potential of colorectal cancer. World J Gastroenterol 13, 4996–5002.

Saigusa, S., Tanaka, K., Toiyama, Y., Matsushita, K., Kawamura, M., Okugawa, Y., Hiro, J., Inoue, Y., Uchida, K., Mohri, Y., et al. (2012). Gene expression profiles of tumor regression grade in locally advanced rectal cancer after neoadjuvant chemoradiotherapy. Oncol Rep 28, 855–861.

Saunders, A., Huang, X., Fidalgo, M., Reimer, M.H., Jr., Faiola, F., Ding, J., Sanchez-Priego, C., Guallar, D., Saenz, C., Li, D., et al. (2017). The SIN3A/HDAC Corepressor Complex Functionally Cooperates with NANOG to Promote Pluripotency. Cell Rep 18, 1713–1726.

Schodel, J., Oikonomopoulos, S., Ragoussis, J., Pugh, C.W., Ratcliffe, P.J., and Mole, D.R. (2011). High-resolution genome-wide mapping of HIF-binding sites by ChIP-seq. Blood 117, e207–217.

Schofield, C.J., and Ratcliffe, P.J. (2004). Oxygen sensing by HIF hydroxylases. Nature reviews. Molecular cell biology 5, 343–354.

Schumm, K., Rocha, S., Caamano, J., and Perkins, N.D. (2006). Regulation of p53 tumour suppressor target gene expression by the p52 NF-kappaB subunit. EMBO J 25, 4820–4832.

Semenza, G.L. (2008). Hypoxia-inducible factor 1 and cancer pathogenesis. IUBMB Life 60, 591–597.

Semenza, G.L. (2009). Regulation of oxygen homeostasis by hypoxia-inducible factor 1. Physiology (Bethesda) 24, 97–106.

Semenza, G.L. (2012). Hypoxia-inducible factors: mediators of cancer progression and targets for cancer therapy. Trends Pharmacol Sci 33, 207–214.

Semenza, G.L. (2013). Cancer-stromal cell interactions mediated by hypoxia-inducible factors promote angiogenesis, lymphangiogenesis, and metastasis. Oncogene 32, 4057–4063.

Semenza, G.L., Jiang, B.H., Leung, S.W., Passantino, R., Concordet, J.P., Maire, P., and Giallongo, A. (1996). Hypoxia response elements in the aldolase A, enolase 1, and lactate dehydrogenase A gene promoters contain essential binding sites for hypoxia-inducible factor 1. J Biol Chem 271, 32529–32537.

Sloan, C.A., Chan, E.T., Davidson, J.M., Malladi, V.S., Strattan, J.S., Hitz, B.C., Gabdank, I., Narayanan, A.K., Ho, M., Lee, B.T., et al. (2016). ENCODE data at the ENCODE portal. Nucleic Acids Res 44, D726–732.

Smith, J.J., Deane, N.G., Wu, F., Merchant, N.B., Zhang, B., Jiang, A., Lu, P., Johnson, J.C., Schmidt, C., Bailey, C.E., et al. (2010). Experimentally derived metastasis gene expression profile predicts recurrence and death in patients with colon cancer. Gastroenterology 138, 958–968.

Smith, K.T., Sardiu, M.E., Martin-Brown, S.A., Seidel, C., Mushegian, A., Egidy, R., Florens, L., Washburn, M.P., and Workman, J.L. (2012). Human family with sequence similarity 60 member A (FAM60A) protein: a new subunit of the Sin3 deacetylase complex. Mol Cell Proteomics 11, 1815–1828.

Statham, A.L., Strbenac, D., Coolen, M.W., Stirzaker, C., Clark, S.J., and Robinson, M.D. (2010). Repitools: an R package for the analysis of enrichment-based epigenomic data. Bioinformatics 26, 1662–1663.

Streubel, G., Fitzpatrick, D.J., Oliviero, G., Scelfo, A., Moran, B., Das, S., Munawar, N., Watson, A., Wynne, K., Negri, G.L., et al. (2017). Fam60a defines a variant Sin3a-Hdac complex in embryonic stem cells required for self-renewal. EMBO J 36, 2216–2232.

Subramanian, A., Tamayo, P., Mootha, V.K., Mukherjee, S., Ebert, B.L., Gillette, M.A., Paulovich, A., Pomeroy, S.L., Golub, T.R., Lander, E.S., et al. (2005). Gene set enrichment analysis: a knowledge-based approach for interpreting genome-wide expression profiles. Proc Natl Acad Sci U S A 102, 15545–15550.

van Uden, P., Kenneth, N.S., and Rocha, S. (2008). Regulation of hypoxia-inducible factor-1alpha by NF-kappaB. Biochem J 412, 477–484.

van Uden, P., Kenneth, N.S., Webster, R., Muller, H.A., Mudie, S., and Rocha, S. (2011). Evolutionary conserved regulation of HIF-1beta by NF-kappaB. PLoS Genet 7, e1001285.

Wada, T., Shimba, S., and Tezuka, M. (2006). Transcriptional regulation of the hypoxia inducible factor-2alpha (HIF-2alpha) gene during adipose differentiation in 3T3-L1 cells. Biol Pharm Bull 29, 49–54.

Wang, G.L., Jiang, B.H., Rue, E.A., and Semenza, G.L. (1995). Hypoxia-inducible factor 1 is a basic-helix-loop-helix-PAS heterodimer regulated by cellular O2 tension. Proc Natl Acad Sci U S A 92, 5510–5514.

Yoshimura, H., Dhar, D.K., Kohno, H., Kubota, H., Fujii, T., Ueda, S., Kinugasa, S., Tachibana, M., and Nagasue, N. (2004). Prognostic impact of hypoxia-inducible factors 1alpha and 2alpha in colorectal cancer patients: correlation with tumor angiogenesis and cyclooxygenase-2 expression. Clin Cancer Res 10, 8554–8560.

Yu, F., White, S.B., Zhao, Q., and Lee, F.S. (2001). HIF-1alpha binding to VHL is regulated by stimulus-sensitive proline hydroxylation. Proc Natl Acad Sci U S A 98, 9630–9635.

Yu, G., Wang, L.G., and He, Q.Y. (2015). ChIPseeker: an R/Bioconductor package for ChIP peak annotation, comparison and visualization. Bioinformatics 31, 2382–2383.

Zhu, L.J., Gazin, C., Lawson, N.D., Pages, H., Lin, S.M., Lapointe, D.S., and Green, M.R. (2010). ChIPpeakAnno: a Bioconductor package to annotate ChIP-seq and ChIP-chip data. BMC Bioinformatics 11, 237.

